# Time-dependent competition between habitual and goal-directed response preparation

**DOI:** 10.1101/201095

**Authors:** Robert M Hardwick, Alexander D Forrence, John W Krakauer, Adrian M Haith

## Abstract

Converging evidence indicates that separate goal-directed and habitual systems compete to control behavior^1^. However, it has proven difficult to reliably induce habitual behavior in human participants^2–4^. We reasoned that habits may be present in the form of habitually prepared responses, but are overridden by goal-directed processes, preventing their overt expression. Here we show that latent habits can be unmasked by limiting the time participants have to respond to a stimulus. Participants trained for 4 days on a visuomotor association task. By continuously varying the time allowed to prepare responses, we found that the probability of expressing a learned habit followed a stereotyped time course, peaking 300-600ms after stimulus presentation. This time course was captured by a computational model of response preparation in which habitual responses are automatically prepared at short latency, but are replaced by goal-directed responses at longer latency. A more extensive period of practice (20 days) led to increased habit expression by reducing the average time of movement initiation. These findings refine our understanding of habits, and show that practice can influence habitual behavior in distinct ways: by promoting habit formation, and by modulating the likelihood of habit expression.

---

Behavioral, computational, and neurobiological evidence suggests that behavior is governed by two distinct systems^5,6^. The goal-directed system selects actions based on an evaluation of which options will best achieve future task goals. In principle, this yields the best possible outcome, but at the cost of being computationally-intensive and slow. By contrast, the habitual system selects actions based on what has been successful in the past. Habitual action selection is simpler and faster, but it is inflexible; it produces the same action irrespective of changes in goals or the environment.

Studies in animals demonstrate that repeating a newly learned behavior leads to a transition from goal-directed to habitual behavior^1,3^. This work has also established dissociable neural substrates for goal-directed and habitual systems, with different regions of the cortex and striatum implicated in expressing goal-directed versus habitual behavior^7^. However, establishing similarly robust habit formation in humans has been surprisingly elusive^2–4^.

We propose that a major reason why habit formation has been difficult to observe in humans is that habits exist, but are masked by goal-directed processes. Actions could be habitually selected and prepared without being immediately executed, allowing them to be replaced by more appropriate, goal-directed responses before any movement is generated. This view is supported by recent work establishing that movement preparation occurs independently from, and much earlier than, movement initiation^8^. Critically, the habitual system provides potential responses rapidly, while the goal-directed system requires longer processing times. We therefore reasoned that limiting the time participants had to prepare responses would prevent preparation of goal-directed responses, unmasking otherwise latent habits.

**Figure 1.**
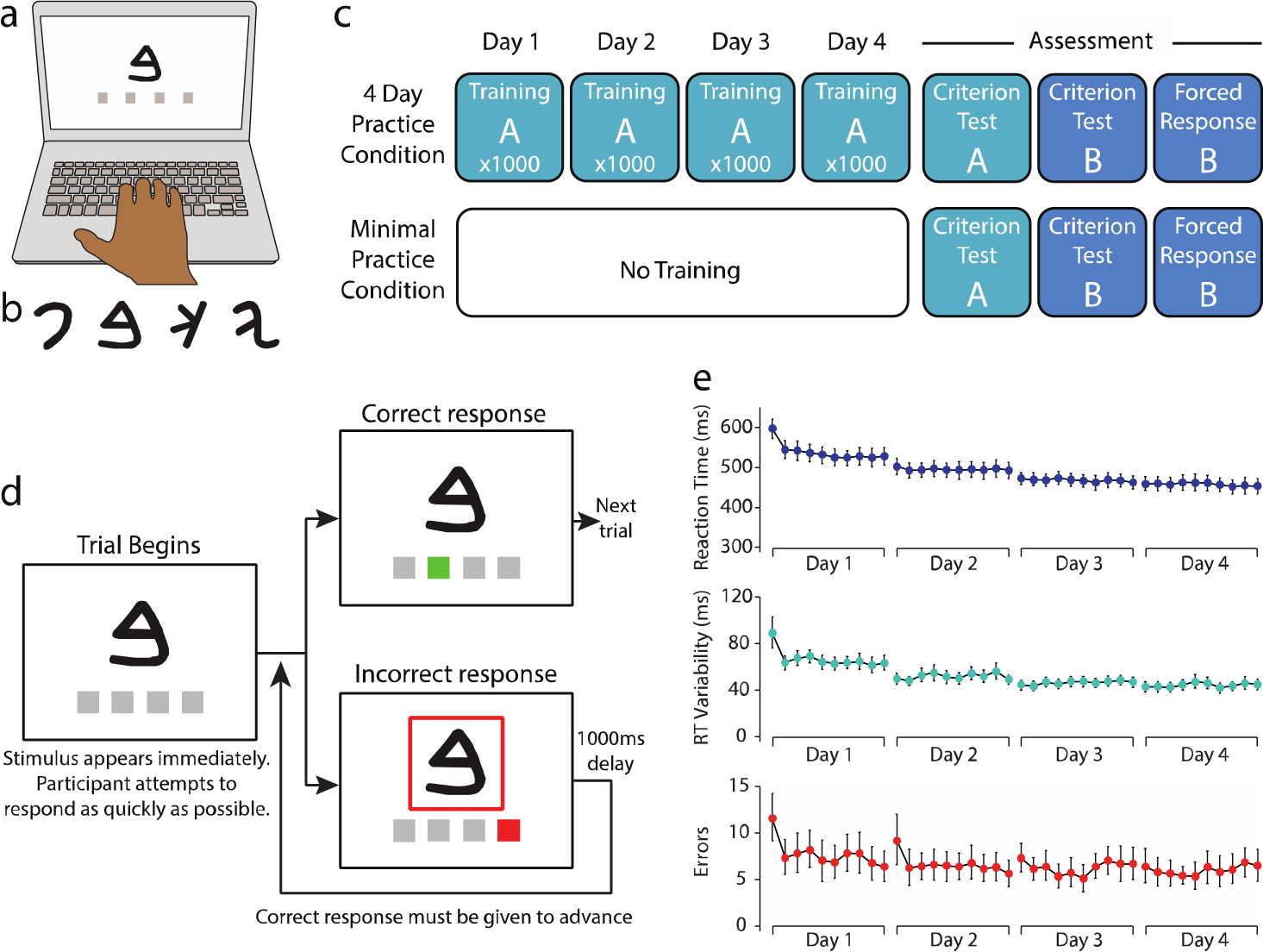
Task and training schedule for Experiment 1. a) Experimental setup. Participants responded to the appearance of a stimulus by pressing a corresponding keyboard button. b) Example stimuli (letters of the Phoenician alphabet). c) Experiment 1 overview. In the 4-Day Practice condition, participants completed 4,000 reaction-time-based training trials on original mapping A over 4 days. On the fifth day (Assessment) they were tested on their knowledge of mapping A in a criterion test block, then learned revised mapping B until they achieved a steady accuracy criterion (see Figure 2d). They then completed forced-response trials (Figure 2e) under this new mapping. In the Minimal Practice condition, participants only performed the assessment session (and therefore practiced the original mapping A only until they achieved a steady accuracy criterion). d) Trial structure of the reaction-time-based training task. Participants attempted to complete blocks of 100 trials as quickly as possible, incurring a 1s time penalty for incorrect responses. e) Data from the reaction-time-based training in the 4-Day Practice condition. Participants’ median reaction times, reaction time variability (median absolute deviation) and error rates improved with training. Error bars represent bootstrapped 95% confidence intervals.

In Experiment 1, participants (n=22) completed a visuomotor association task in which arbitrary stimuli directed them to press specific keyboard buttons (Figure 1a). We contrasted behavior in two conditions: a *4-Day Practice* condition and a *Minimal Practice* condition. In the 4-Day Practice condition, participants first trained on a previously unseen stimulus-response mapping, completing 4,000 reaction-time-based trials (100 trials × 10 blocks × 4 days) in which they responded as quickly as possible to visual stimuli presented in rapid succession (Figure 1d). Performance at this task improved with practice (Figure 1e), with significant reductions in reaction times (average of first vs last day, t-test, *t*_21_=11.96, *p*<0.001), reaction time variability (t-test on reaction time median absolute deviation, *t*_21_=9.38, *p*<0.001), and errors (t-test, *t*_21_=2.18, *p*<0.05).

We assessed whether practice led participants to habitually prepare a particular response by transposing the required responses for two of the four stimuli (Figure 2b). Participants first learned this revised mapping in a criterion test block; they were instructed that there were no time constraints in this block, and that they should focus on learning a new stimulus-response mapping. They trained on the revised mapping until satisfying an accuracy criterion of five consecutive correct responses to each stimulus, which occurred on average within 44±5 (mean+SEM) trials. The number of trials needed was comparable to that required to learn the original mapping at the start of the Minimal Practice condition (40±4 trials) (Figure 2d; paired samples t-test, *t*_21_=0.64, *p*=0.53). Thus, participants had no difficulty in learning to accurately respond according to the revised mapping, regardless of whether they had practiced an incompatible mapping beforehand. This seemingly suggested that four days of practice had not led the association to become habitual.

**Figure 2.**
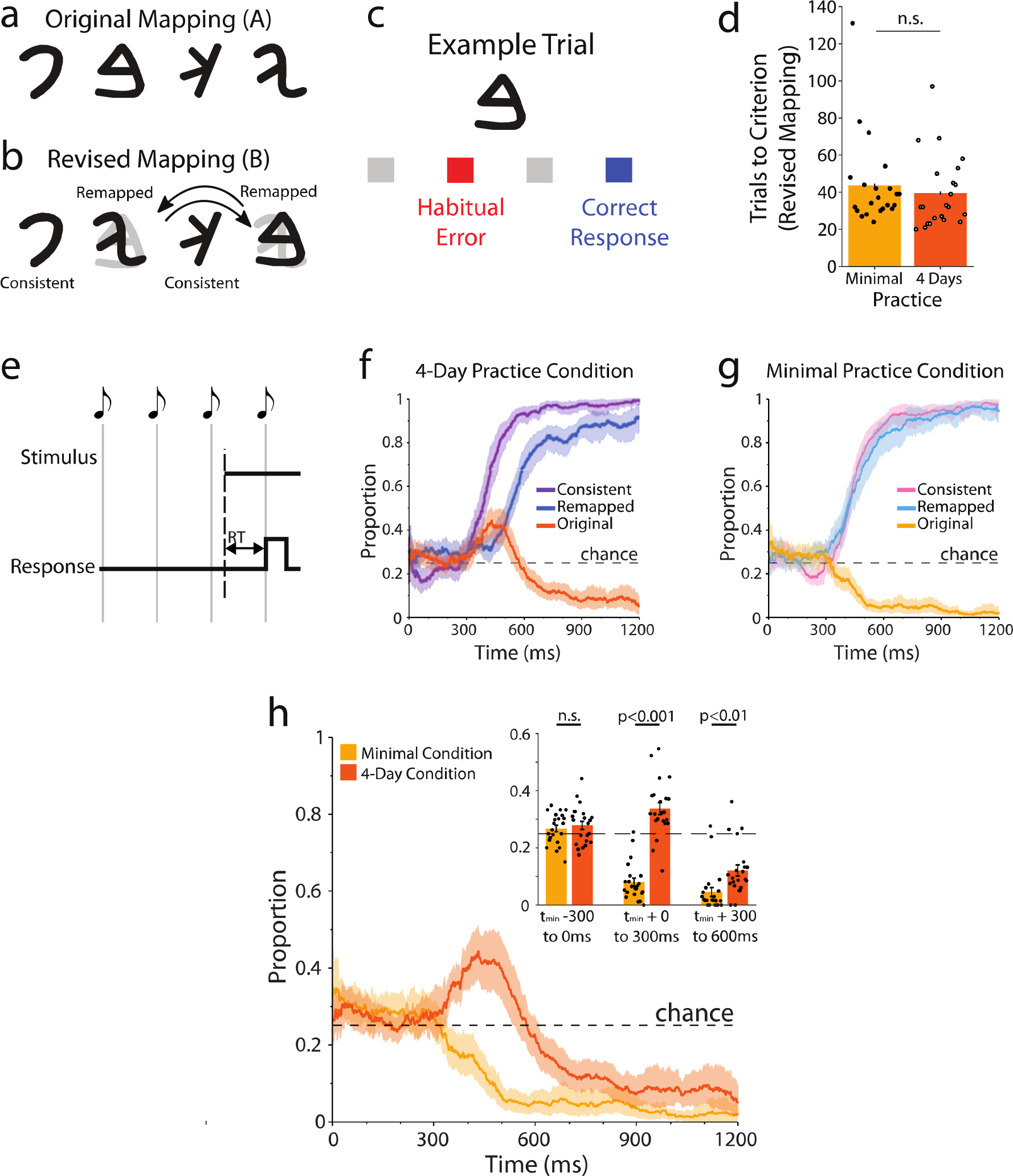
Switch manipulation, retraining, and forced-response task results. a) Following either minimal practice or four days of practice on original stimulus-response mapping A (example mapping shown), two stimulus-response associations were switched to create a revised mapping (b). c) This revised mapping allowed the identification of habitual behavior - trials in which participants acted according to the originally learned mapping (a habitual error) rather than the revised mapping (correct response). d) Participants trained on the revised mapping without reaction time constraints until they reached a criterion of 5 consecutive correct responses to each stimulus. Participants required approximately 40 trials to learn this revised mapping, regardless of the volume of training they had completed on the original mapping. Error bars represent ±1SEM. e) Participants then completed a forced-response task, shown here in schematic form. Participants were instructed to respond synchronously with the final tone in a sequence of four equally spaced tones. Stimulus onset was varied relative to this time in order to impose differing response times, distributed randomly and uniformly from 0-1200ms prior to the fourth tone. f) Distribution of responses as a function of allowed preparation time for the 4-Day Practice group. Each line presents the proportion of different types of responses within a 100ms sliding-window. Purple: correct responses for unchanged, consistently mapped stimuli. Blue: Correct responses for remapped stimuli. Red: habitual errors for remapped stimuli. The brief increase in the probability of responding according to the original map between 300-600ms indicates transient habitual preparation of the originally practiced mapping. g) Analogous results for the Minimal Practice condition, indicating no evidence of habitual action preparation. h) Direct comparison of the time-varying probability of expressing the original response across the groups. Inset compares the proportion of habitual responses binned across successive 300ms intervals relative to the minimum time at which participants could respond to stimuli (t_min_ - see text for details). Shaded error regions in f-h represent bootstrapped 95% confidence intervals. Bar chart error bars represent ±1SEM.

We then examined whether limiting response preparation time would unmask habitual preparation of the initially practiced response. We achieved this using a “forced-response” paradigm^8–10^ (see methods). Participants were forced to initiate a response at a fixed time in each trial (i.e. synchronously with the fourth tone of a metronome). Varying the onset of the visual stimulus in relation to the time of the fourth tone enabled us to control the time the participants had to prepare their action (Figure 2e), allowing us to probe which responses were prepared at different times.

To visualize the time-course of response preparation, we assessed the probability of expressing different responses within a 100ms sliding window (Figure 2f). Focusing first on responses to stimuli that did not change between the mappings (consistently mapped stimuli; purple curve in Figure 2f), the probability of generating the correct response was at chance (0.25) for preparation times less than ≈300ms. This indicates that participants had insufficient time to process the stimulus, and therefore had to guess. The probability of generating the correct response then rose gradually as preparation time increased, reaching asymptote between 700-900ms.

Habitual response preparation was revealed by assessing the time course of responses to remapped stimuli. In the 4-Day practice condition, the probability of generating the originally learned response (Figure 2f, orange curve) began at chance for low (<300ms) preparation times but then transiently increased above chance at preparation times of 300-600ms, before declining towards zero as the proportion of correct responses (Figure 2f, blue curve) began to increase. Therefore, despite the fact that participants had no difficulty in acquiring the revised mapping after four days of practice, forcing them to respond at low preparation times unmasked latent habitual responses. By contrast, in the Minimal Practice condition (Figure 2g), the proportion of habitual errors (Figure 2g, yellow curve) began at chance, then declined as preparation time increased.

We summarized and confirmed these observations by analyzing the overall likelihood of habitual responses in a 300ms interval aligned to the minimum possible time at which participants could generate an accurate response (t_min_, identified as the time at which a cumulative Gaussian fit to the speed-accuracy trade-off for consistently mapped stimuli first reached 5% of its height), which showed a significant interaction between condition and required preparation time (_RM_ANOVA, F_1,21_=58.32, p<0.001).

We note that participants’ ability to prepare appropriate responses was slowed; the speed-accuracy trade-off for remapped stimuli was shifted later relative to that for consistently mapped stimuli (Figure 2f; purple vs blue curves, significant difference between the center of the speed-accuracy trade-offs, t_21_=5.93, p<0.001, mean difference 93ms). In the Minimal Practice Condition, by contrast, there was no such difference in the speed of preparation between remapped and consistently mapped stimuli (t_21_=0.64, p=0.53).

The time-varying expression of habitual or goal-directed behavior was accurately described by a computational model of response preparation (Figure 3). We first considered a model of response preparation in the case of a single learned association^8^ (Figure 3a). The time at which responses became prepared, *T*_*A*_, was assumed from trial to trial according to a Gaussian distribution. Critically, the response is not necessarily initiated at time *T*_*A*_, but is instead held in a prepared state until a response is initiated. If the time at which the response is initiated is equal to or greater than *T*_*A*_, the participant will have had sufficient time to process the stimulus and prepare the appropriate response. But if the time allowed to respond is less than *T*_*A*_, the participant will not have had enough time to process the stimulus, and will therefore respond at a chance level of accuracy. These assumptions predict a speed-accuracy trade-off qualitatively matching that observed for the consistently mapped stimuli (Figure 3a, lower panel). Improvements in the speed-accuracy trade-off through practice are explained by improvements in the mean and variance of the distribution of *T*_*A*_(Figure 3b, upper panel).

**Figure 3.**
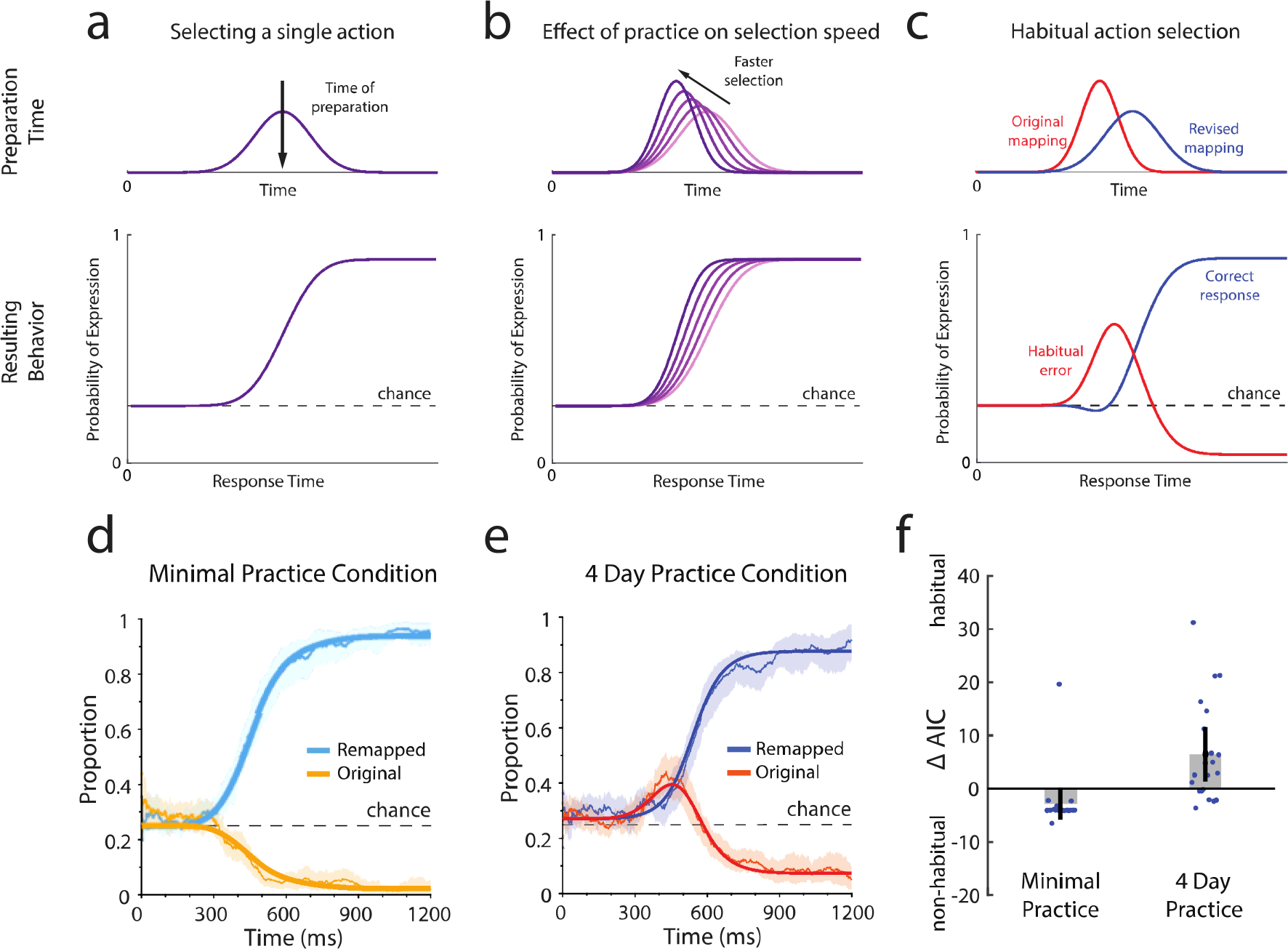
Computational model of response preparation. a) We assume that, in each trial, a response is prepared at a random time (assumed here to follow a Gaussian distribution) after stimulus onset (a, upper panel), giving rise to the observed speed-accuracy trade-off across trials (a, lower panel). b) As the mean and variance of the time of preparation improve (b, upper panel) the speed-accuracy trade-off improves (b, lower panel). c) After the mapping is revised, the original mapping (red) is still habitually prepared. The appropriate, revised response (blue) becomes available later and replaces the habitually prepared response. This sequence of events leads to a time-varying probability of each response being expressed (c, lower panel). d) Averaged behavior (thin lines) and average model fit (bold lines) for the minimal practice condition. Fit shown here for the model with no habitual preparation (ρ = 0; equivalent to panel a), which had a lower AIC for this condition. Error bars show +/− SEM. e) Behavior and model fit in the 4-Day practice condition. Fit shown is for the model with habitual preparation (panel c), which had a lower AIC for this condition. f) Difference in AIC between the habitual (ρ = 1) and non-habitual (ρ = 0) models for individual participant fits to the data. Results favored the habitual preparation model in the majority of participants in the 4-Day Practice condition, but in only one participant in the Minimal Practice condition.

We extended this model to account for the possibility of multiple, competing response-selection processes (Figure 3c). In this model, habitual and goal-directed processes select potential responses in parallel, but compete for preparation of a single action. We assume that the habitual response becomes available at time *T*_*A*_, while the correct, remapped response becomes available at a later time *T*_*B*_. We assume that *T*_*A*_ and *T*_*B*_ are both Gaussian distributed and independent, but with *T*_*B*_ having a greater mean and variance than *T*_*A*_. Critically, participants will prepare the habitual response as soon as it is available (i.e. at time *T*_*A*_), but this will be replaced by the goal-directed response once that becomes available (i.e. at time *T*_*B*_). Participants will express whichever response is prepared at the time of movement initiation. Therefore a habitual response will be generated if the response is initiated after *T*_*A*_ but before *T*_*A*_. This model replicated the stereotypical time course we observed experimentally (Figure 3c, lower panel).

Applying this model, we first compared fits to data pooled across participants. The habitual preparation model accounted for the data in the 4-Day Practice condition significantly better than an alternative model in which the previously practiced response was never prepared (ΔAIC in favour of habitual preparation model = 275.6). By contrast, data from the Minimal Practice condition was best explained by the model in which participants did not habitually prepare actions (ΔAIC in favour of no habitual preparation model = 215.3).

Our models also accounted for individual participant behavior well (Figures S1 and S2). In the 4-Day practice condition, the habit model better explained data for 16/22 participants (mean ΔAIC = 11.3, Figure 3f) suggesting that the majority of participants had become habitual following practice. In the Minimal practice condition, the no-habitual-preparation model better explained behavior in 21/22 participants (mean ΔAIC = 14.8, Figure 3f). Thus, the results of Experiment 1, along with the computational model, established that four days of practice led to habitual behavior in the majority of participants. This habit was not apparent when participants completed the task under self-paced conditions (i.e. in the criterion test blocks), but was unmasked by forcing them to respond rapidly.

In a second experiment we investigated the effects of more extensive practice on habitual behavior. In Experiment 2 a new group of participants (n=14) trained on an original mapping over a period of 4 weeks, completing 20 days of practice in total (1,000 trials per day, 20,000 total trials). As in Experiment 1, participants trained in reaction-time-based trials. The prolonged practice enabled participants to reduce their reaction times by 70ms relative to participants at the of the 4-Day practice condition (group-by-day interaction for first/last day comparison, F_1,34_=22.53, p<0.001; no baseline difference between groups on day 1 of training, t_34_=0.89, p=0.38). The speed of participants’ response preparation was also periodically assessed during learning through forced-response trials, yielding speed-accuracy trade-offs for the practiced mapping (Figure 4). This data revealed a clear improvement in the latency at which participants could generate an accurate response (rmANOVA on mean preparation time, F_4,52_=41.81, p<0.001; Figure 4b).

**Figure 4.**
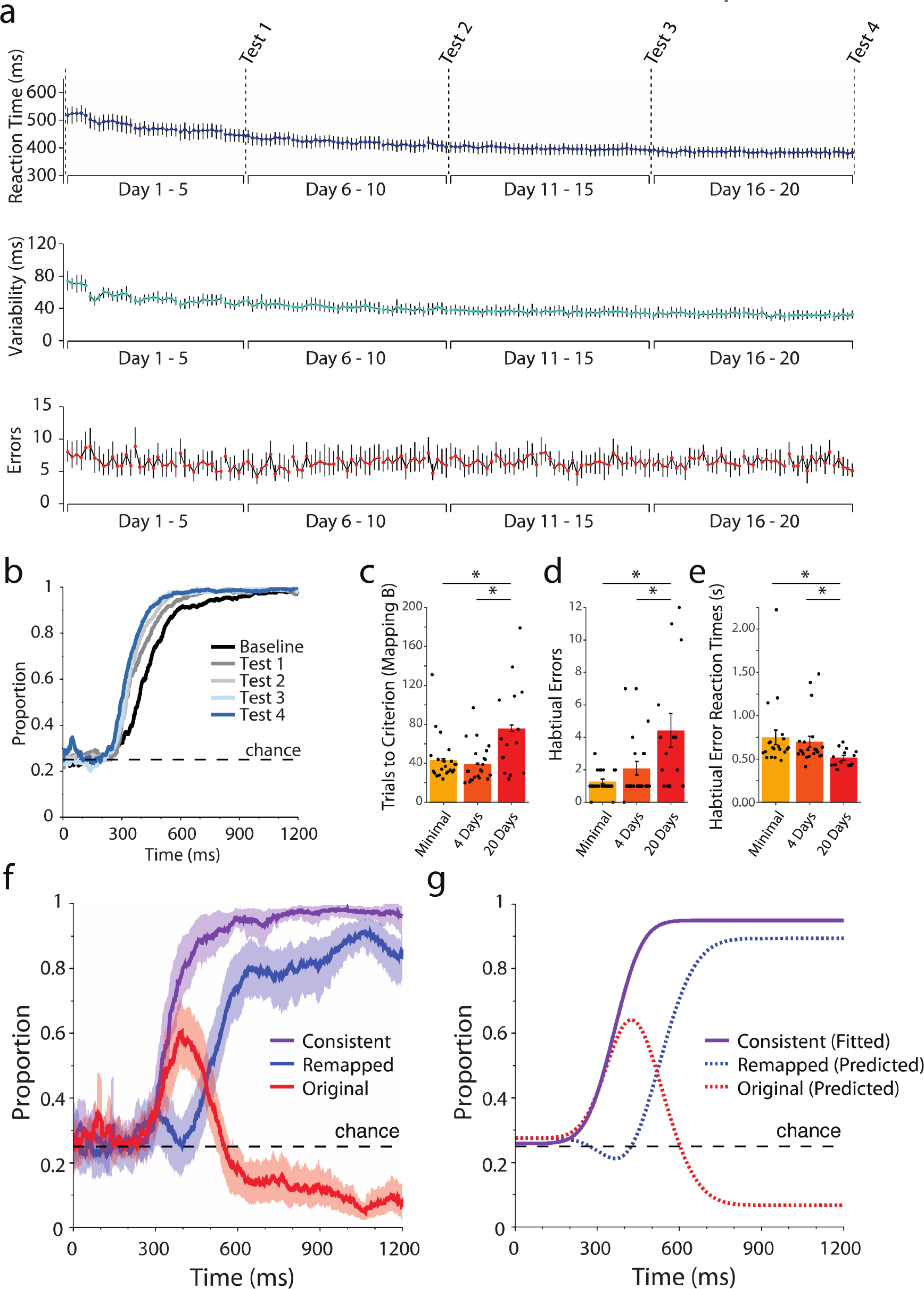
Results for Experiment 2 (20 Days/20,000 trials of Practice). a) Participants’ median reaction times, reaction time variability (median absolute deviation) and error rates over training. Each circle represents data for one block of 100 trials. Separations in the lines between points indicate between-day breaks. Error bars present bootstrapped 95% confidence intervals. b) The speed-accuracy trade-off for the original stimulus-response associations (Mapping A) was assessed using forced-response trials at multiple points during training: at baseline (just after achieving criterion), and at the end of each week of practice. The leftward shift indicates that participants learned to execute the association more rapidly with practice. c) The number of trials required to achieve criterion for the revised mapping was higher in the 20-Day Practice condition than the Minimal Practice or 4-Day Practice conditions in Experiment 1. d) Participants in the 20-Day Practice condition made significantly more habitual errors during the criterion test block under the new mapping, but did so with significantly faster reaction times (e) compared to conditions with less practice. f) Forced-response data for the 20-Day Practice condition. Speed-accuracy trade-off for consistently-mapped stimuli (purple), and remapped stimuli (blue). The peak probability of expressing the originally practiced response (red) was even greater than in the 4-Day Practice condition (see text for statistical comparisons). g) Predicted behavior in the 20-Day Practice condition (dotted lines), based on group-level model fits to behavior in the 4-Day Practice condition, and fits to the speed of preparation of responses to consistently mapped stimuli (solid line).

Following practice, we tested whether response preparation had become habitual by remapping two of the four stimuli, as in Experiment 1. Participants practiced this new mapping until they made 5 consecutive correct responses to each stimulus. Participants needed an average of 76±13 trials to achieve this criterion; significantly more than the 44±5 trials needed in the Minimal Practice condition (t-test, t_34_=2.74, p<0.05) or the 40±4 trials needed in the 4-Day practice condition (t-test, t_34_=3.24, p<0.01). This increase was largely attributable to participants committing more habitual errors (Figure 4d, Mann-Whitney U test vs Minimal Practice condition, Z=2.92, p<0.01, and vs 4-Day Practice condition, Z=2.14, p<0.05).

We also found that participants who learned the revised mapping after 20 days of practice had shorter reaction times during the criterion test block than those who practiced for only four days (Figure 4e, Mann-Whitney U test vs 4-Day Practice condition, Z=2.46, p<0.05), or those that had minimal practice (vs Minimal Practice condition, Z=2.94, p<0.01), suggesting a link between habitual responses and short response times.

Participants were then tested under forced-response conditions (Figure 4f), allowing more detailed examination of the effect of additional practice on their response preparation. As expected from the practice-based improvements in reaction time (Figure 4a) and the speed-accuracy trade-off (Figure 4b), the center of the speed-accuracy trade-off for consistently mapped stimuli (Figure 4f, purple line) was significantly earlier than that of participants in the 4-Day practice condition in Experiment 1 (t-test on mean preparation time, t_34_=2.32, p<0.05, mean difference 48ms).

The speed at which responses to remapped stimuli could be prepared (Figure 4f, blue line) was slower than for consistently mapped stimuli (t-test on mean preparation time, t_34_=11.50, p<0.001, mean difference 162ms), but was similar to that observed after 4 days of practice (t-test on revised response speed-accuracy trade-offs, t_34_=0.81, p=0.43). Thus, given sufficient time, participants were perfectly capable of producing accurate goal-directed responses.

As expected, forced-response trials revealed habitual response preparation in the 20-Day Practice condition. Responses were similar to those in the 4-Day Practice condition, except that participants who trained for 20 days were more likely to produce habitual responses when forced to respond at low latencies (t-test on 20-Day practice vs 4-Day practice groups for 0-300ms following t_min_, t_34_=2.98, p<0.01).

As with the 4-Day Practice condition, data from the 20-Day Practice group favored the habitual preparation model over the no-habitual-preparation model, even more strongly than for Experiment 1 (ΔAIC in favour of habitual preparation model = 318.2). At the individual-participant level, the data favored the habit model in all 14 participants who completed the experiment (mean ΔAIC = 24.3; Figure S3).

Our data indicated that 20 days of practice led to more pronounced habitual behavior compared to just 4 days of practice, which was apparent in two distinct ways. First, participants made more habitual errors in the criterion test block. While this could suggest that the habit had become more difficult to override with goal-directed responses, data from the timed-response condition did not support this view; the speed-accuracy trade-off for generating an accurate, goal-directed response (blue lines in Figure 2f versus Figure 4f) was indistinguishable from that in the 4-Day practice condition. Instead, the increase in the number of habitual errors occurring during this block was more likely attributable to the fact that participants who trained for 20 days responded with lower reaction times during the criterion test block. This suggests participants may have persisted in responding with short reaction times that were successful during their training^11^, which in turn increased the likelihood that they would express the habitually prepared action.

The model predicted the fact that the peak probability of expressing a habitual error was greater after 20 days of practice than after 4 days of practice (Figure 4g). The probability of expressing a habitual error is strongly influenced by the speed at which responses can be prepared; more rapid preparation of a habitual response broadens the window of opportunity for it to be expressed before it is replaced by the appropriate, goal-directed response (Figure S4a). After 20 days of practice, participants had significantly reduced the latency at which they could select and prepare the habitual response (Figure 4b). The increase in the peak probability of a habitual response observed between 4 and 20 days of practice was consistent with model predictions based on observed improvements in preparation speed (see Figure 4f; see Supplemental Materials for further details).

An alternative interpretation of the increased probability of generating a habitual error is that habitual preparation became more likely to occur. Our initial model considered habitual response preparation as an all-or-nothing process – a habitual response is always prepared when available (i.e. when response time is greater than *T*_*A*_ and less than *T*_*B*_). However, we also considered a model in which habitual preparation of the originally practiced mapping might occur with a variable probability *ρ*_*A*_, which could be construed as the ‘strength’ of the habit. Under this model, with a strong habit (*ρ*_*A*_ = 0), the originally practiced mapping would be reliably prepared every time a particular stimulus is presented. With no habit (*ρ*_*A*_ = 0), the originally practiced mapping is never prepared. With an intermediate habit (e.g. 0 < *ρ*_*A*_ < 1) the originally practiced mapping might be prepared on only a subset of trials. Practice may increase the probability of habitual response preparation occurring at all in a given trial, i.e. strengthening the habit. We fit this model to pooled data at the group level, finding that this probabilistic habit model better explained the data than the original habit model for both the 4-day condition (ΔAIC = 10.4) and the 20-day condition (ΔAIC = 11.7). However, we noted that this superior fit at the group level may not have been due to individual participants probabilistically preparing habitual responses (i.e. engaging in habitual preparation only in a subset of trials), but could instead be attributable to the group data containing a mixture of habitual and non-habitual participants. Fits to individual participants showed that the probabilistic habit model (*ρ*_*A*_ ∊ [0,1]) better accounted for the 4-day practice condition data in only 4/22 participants, compared to 12/22 for the original habit model (*ρ*_*A*_ = 1), and 6/22 for the no-habit model (*ρ*_*A*_ = 0). In the 20-day practice group, the original habit model (*ρ*_*A*_ = 1) best explained the data for all 14 participants. Our results therefore offered strongest support for the interpretation that habit formation is a discrete, all-or-nothing process.

In summary, Experiment 2 illustrates that extensive practice led to more overt habitual behavior. This effect occurred because well-practiced participants tended to respond at shorter latencies and could prepare the habitual response more rapidly, rather than habitual preparation itself becoming more likely to occur in any given trial.

The relative expression of different components of learning has previously been shown to be influenced by limiting cognitive resources^12,13^, including the time available to prepare an action^14–18^. However, previous research has manipulated preparation time in a relatively simple ‘high-or-low’ manner, or based on spontaneous variations in ‘voluntarily’ selected reaction times. The forced-response paradigm used here allowed us to systematically assess the temporal dynamics of these effects. Examining responses across a continuum allowed us to precisely track the evolving competition between habitual and goal-directed response selection.

Our results and model clarify the nature of competition between habitual and goal-directed response selection. Although both processes can select specific actions in parallel, they compete for which of these potential responses becomes prepared and ready to execute. This is consistent with the emerging view that, although multiple goals might be entertained in parallel, only a single movement is ever prepared^19,20^.

Behavioral and neuroscientific evidence indicates that the preparation and initiation of actions occur separately^8,21^. Our results and model support this view, indicating that a response can be habitually selected and prepared, but not immediately expressed, allowing it to later be replaced by a goal-directed response. Importantly, these preparatory events do not directly influence the timing of response initiation, which is thought to be under independent control^8^ and subject to its own use-dependent biases^11^. Critically, when participants respond with suitably low latencies, they will generate a habitual response. This accounts for the differences in the behavior in criterion test blocks following training. In Experiment 1 participants self-selected relatively long response times; consequently, no habitual responses were apparent. By contrast, in Experiment 2; participants appeared to persist with the low reaction times that had been successful during training blocks, and were therefore more liable to express their habit.

In both experiments, practice enabled participants to generate accurate responses at lower latencies, as reflected in an improved speed-accuracy trade-off (i.e. they became more skilled at the task^22–24^). Both skilled and habitual behavior are hallmarks of automatic performance^25^. Although definitions of automaticity vary, it is typically considered to involve improvements in skill, habitual behavior, and the ability to perform actions with little or no conscious attention^25–27^. Notably, our data indicate that skill improved continuously with practice, while the presence of habitual action preparation was better explained as an all-or-nothing phenomenon (see supplementary materials), suggesting a potential dissociation between the processes of skill acquisition and habit formation in automatic behaviour.

Previous research has failed to achieve any clear consensus on the neural basis of automaticity, proposing that it arises either through increases in network efficiency^26,27^, or through discrete shifts in the brain regions that control behavior, either within the basal ganglia, within cortex, or from the cortex to the cerebellum^28–30^ (note that similar regions are also frequently implicated in both skill acquisition^31,32^ and habit formation^6^). We propose these differing conclusions may have arisen because the tasks used to examine automaticity likely engaged multiple distinct learning processes: skill learning, habit formation, and changes in mental representation, and may have engaged these to differing degrees. Employing separate measures of skill acquisition and habit formation could therefore significantly help to clarify the underlying neural basis of automaticity.

In summary, our present results establish the existence of habitual response preparation, and demonstrate how fine-grained behavioral assessment can unmask habits that may not be apparent under conventional, self paced approaches. Practice led to the formation of a habit within four days. However, further practice also brought about additional changes that made an existing habit more likely to be expressed. Thus, our results highlight an important distinction between formation and expression of habits. We suggest that dissociating these aspects of habitual responding is critical to achieve a complete understanding of habitual behavior.

## Methods

### Participants

A total 39 participants took part in the study. Experiment 1 included 24 individuals. Two participants withdrew from Experiment 1 having completed only one of the two required experimental conditions, leaving 22 full datasets for the experiment (17 right handed, 13 female, mean age 21 years). Experiment 2 included 15 participants. One participant withdrew due to computer hardware failure, leaving a total of 14 participants (12 right handed, 4 female, mean age 26 years) to complete the experiment. All participants gave written informed consent, and all procedures were approved by the Johns Hopkins School of Medicine Institutional Review Board. Participants received financial compensation ($15/hour) for their participation.

### General Procedures

The task involved responding to the appearance of one of four stimuli (letters of the Phoenician alphabet) by pushing a specific key on a computer keyboard with the index, middle, ring, or little finger of the dominant hand. The stimulus corresponding to each response was counterbalanced across participants, controlling for potential effects whereby participants would find some stimuli easier to recognize and learn to respond to than others. As Experiment 1 comprised two conditions and used a within-subjects design, we employed two sets of distinct stimuli (see Figure S5), and counterbalanced the condition to which they corresponded across participants. Participants in Experiment 1 also completed the two conditions (Minimal Practice and 4-Day Practice) in a counterbalanced order. In all conditions stimuli were presented psuedorandomly within 20 trial subblocks, with each stimulus appearing five times within a subblock, and the same stimulus appearing in two consecutive trials at the most. Participants attempted to respond to stimuli in *training, criterion test,* or *forced-response* trial blocks:

### Training blocks

During training, participants completed a gamified task in which they attempted to complete blocks of 100 reaction-time based trials as quickly as possible (See Figure 1d). In each trial a stimulus appeared in the center of the screen, and a tone played to signal the participant that a trial had started. On correct responses a pleasant auditory tone sounded, and after a 300ms delay the task advanced to next trial. Errors were punished with an auditory buzzer sound and a compulsory delay of 1000ms, after which the participant could once again respond to the same stimulus; this process repeated until the correct response was provided, at which point the task progressed to the next trial. At the end of each block participants received feedback on the time taken to complete each block, and how this compared to their ‘personal best’ block completion time. Participants were encouraged to improve their performance by aiming to beat their personal best time each time they completed the task.

### Criterion test of mapping knowledge

We assessed the ability of participants to learn new or revised stimulus-response associations using criterion test blocks. Participants were instructed that reaction time constraints were removed, that their goal was to learn the correct set of stimulus-response associations, and that the block would end once they had made enough correct responses in a row. These blocks ended once participants had made five consecutive correct responses to each stimulus (minimum of 20 trials), and the number of trials required to reach this steady, high-accuracy criterion was recorded.

### Forced-response blocks

We used forced-response trials to probe the speed of response preparation and to assess whether participants habitually prepared their responses. Each block comprised 100 trials. In each trial the participant heard a series of four tones, spaced 400ms apart, and was instructed to synchronize their response with the onset of the fourth and final tone. The stimulus appeared at a random (uniformly distributed) time during the series of tones, effectively controlling the amount of time in which the participant could prepare their response. As such, in cases in which participants did not have chance to process the stimulus (i.e. when it appeared less than ~300ms before the deadline of the fourth tone), they were essentially forced to guess the correct response (and thus had a 1 in 4 chance of responding correctly).

## Protocol

### Experiment 1

In Experiment 1 participants completed a counterbalanced, crossover design comprising two conditions. Both conditions began with a warm-up/familiarization task. Participants completed 2 blocks (200 trials total) of reaction-time based trials in response to non-arbitrary stimuli (pictures of the hand with one finger colored black to indicate the desired response - see Figure S6). This was followed by 2 blocks (200 trials total) of forced-response trials to the same non-arbitrary stimuli. This familiarization period allowed the experimenter to explain the practice and forced-response paradigms to the participant, and to ensure that the participant could comply with the demands of each task.

Following this familiarization procedure, participants in the Minimal Practice condition then learned an original map of stimuli (Mapping A) in a block of criterion test trials, after which a second block of criterion test trials was used to introduce and assess the ability to learn a revised mapping (mapping B). We then probed for habitual response preparation using forced-response trials. The 4-Day Practice condition used the same assessment, but this was completed after four consecutive days of practice (10×100 trial reaction time training blocks each day) on Mapping A.

### Experiment 2

The second experiment comprised a single condition. All participants first completed the same familiarization procedure as in Experiment 1. Participants then completed a criterion test block in which they learned a set of stimulus-response associations through trial and error (Mapping A). Once they had achieved this criterion, they completed 500 forced-response trials on this original mapping (to allow assessment of baseline performance), followed by 500 reaction-time-based training trials. Each day thereafter, participants completed ‘training sessions’ in which they completed 10×100 trial blocks of reaction-time-based training trials. On the final (fifth) day of training for each week of practice, participants completed a ‘training and probe’ session, in which they completed 500 (5 x 100-trial blocks) reaction-time based training trials, followed by 500 (5×100 trial blocks) of forced-response trials. Participants completed 20 sessions in this manner (aiming to complete five sessions of practice in each seven-day week), allowing us to measure changes in performance at baseline, and after each of the four weeks of practice.

On a separate day after all training sessions were complete, participants were exposed to the same assessment as in Experiment 1; they learned a revised set of stimulus-response associations in a criterion test block, and their performance on this new mapping was then probed in 5×100 trial blocks of forced-response trials.

## Data Analysis

### Reaction time trials

Performance for each block was measured by taking the median reaction time (measured from stimulus onset to response onset) for correct trials, the median absolute deviation of the reaction time, and by calculating the error rate for each block (i.e.number of erroneous responses in each block; note that it was possible for participants to make multiple errors in the same trial, as the trial did not advance until the participant provided the correct answer).

### Criterion test trials

Criterion test trials were primarily analyzed by counting the number of trials required for a participant to make five consecutive correct responses to each stimulus. The reaction time and accuracy for each response was recorded (although participants were made aware that there were no reaction time requirements for these trials).

### Forced-response trials

Preparation times were calculated as the time between stimulus presentation and the participant’s first response. We examined the probability of three types of response; correct responses to *consistently mapped* stimuli,( i.e. stimuli for which the same key press was required throughout the experiment), correct responses to the *remapped* associations, and responses consistent with the *original* mapping. A sliding window visualized the time-varying probability for each of these trial types and response types; responses were binned over 100ms windows, and the proportion of correct vs total responses was calculated and recorded for the center of each window.

## Response Preparation Models

First, we consider the simple case of generating responses in the context of a single mapping *A* between stimuli and actions (see^8^). We assume that preparation of the response (i.e. becoming ready to generate the response) occurred as a discrete event at a random time 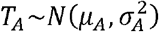. Responses generated prior to *T*_*A*_ are assumed to be roughly uniform across the four possible keys, reflecting the fact that participants were guessing. Responses generated later than *T*_*A*_ are assumed to be correct with probability *q*_*A*_ (with other responses distributed uniformly across the other three keys). The probability of observing a correct response, given that the response was generated at time *t* being correct is therefor given by

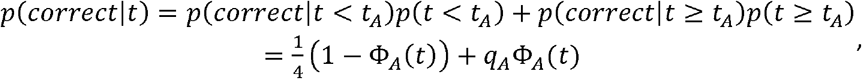

Where 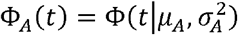 is the cumulative distribution of *T*_*A*_. The probability of an error is, likewise given by

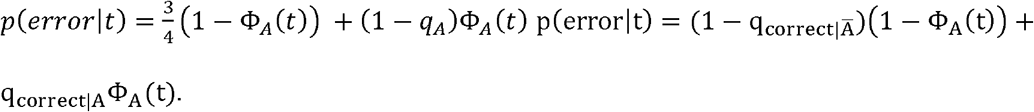

We extended this model to include the possibility of two distinct mappings A (the original, habitual mapping) and B (the goal-directed mapping). We assumed that the associated responses become available at independent times 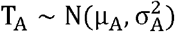 and 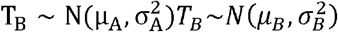 with *T*_*B*_ > *T*_*A*_, reflecting the fact that we expect response B to be available, on average, later than response A. In this instance, when the presented symbol is one that has been remapped, there are three possible response types: 1) a correct response (according to the ‘true’ mapping B), 2) a habitual error (correct according to the original mapping, A), and 3) an “other error” (not correct for either A or B). The probability of each response type depends on which of the events, A or B, have occurred by the time of responding. To simplify notation, we use *Ā* to denote the event *t* < *T*_*A*_, and *A* to denote the event *t* ≥ *T*_*A*_ with likewise definitions for *B̅* and *B*. The probability of a given response being generated at time *t* is then given by:

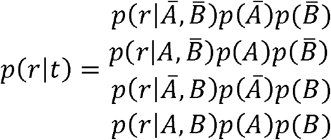

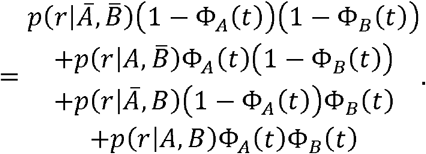

We assumed that the habitual and goal-directed processes select the response associated with that mapping with probability *q*_*A*_ and *q*_*B*_, respectively. Critically, we assumed that when the goal-directed response B becomes available it will immediately be prepared, displacing any habitually prepared response A. Additionally, the likelihood of different responses might have been influenced by a possible bias in guessing either remapped or non-remapped keys, which we represented through the parameter *q*_*I*_, reflecting the baseline probability of pressing one of the two remapped keys. The overall distribution of responses is then given by:

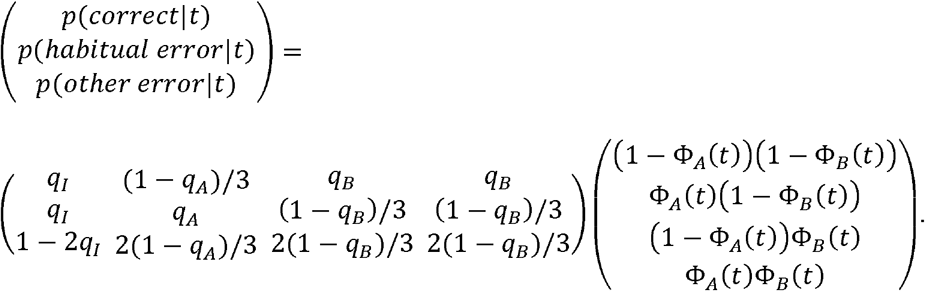

This model can be more compactly expressed as:

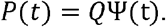

Finally, we extended this model to allow for the possibility that event A or B occurring failed to update the prepared response. We assumed success rates *ρ*_*A*_and *ρ*_*B*_ respectively for these events. Note that these parameters are distinct from *q*_*A*_ and *q*_*B*_, which represent the probability of updating the prepared response to the appropriate key. The parameter *ρ*_*B*_ allows the model to capture the possibility of a persistent propensity to committing habitual errors even at long reaction times, due to a lapse in generating the goal-directed response. The parameter *p*_*A*_ can be interpreted as a ‘habit strength’ parameter, governing the probability of habitual response preparation occurring. Mathematically, the probability of each response type, allowing for potential lapses, is given by:

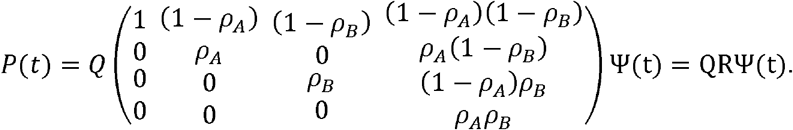

This full model included 9 parameters in total. However, *q*_*A*_ and *ρ*_*A*_ led to qualitatively similar effects on behavior. We therefore set *q*_*A*_ = 0.95, a nominal value reflecting typical success rates for already well-known mappings.

### Model variants

We refer to the full model described above, with 8 free parameters (*μ*_*A*_,*σ*_*A*_,*μ*_*B*_,*σ*_*B*_,*q*_*A*_,*q*_*B*_,*ρ*_*A*_*ρ*_*B*_), as the *probabilistic habit* model. Two further models emerge as special cases: the *habit* model corresponds to *ρ*_*A*_ = 1, i.e. habitual preparation on every trial, having 7 parameters. The *no-habit* model corresponds to *ρ*_*A*_ = 0. In this model, the parameters *μ*_*A*_,*σ*_*A*_ have no effect on behavior and *q*_*B*_ and *p*_*B*_ become equivalent. The no-habit model therefore has four free parameters:*μ*_*B*_,*σ*_*B*_,*q*_*A*_,*q*_*B*_.

We estimated the parameters of these models based on maximum likelihood, using the Matlab function fmincon. To achieve more robust fits to data that avoided the possibility of unrealistically steep speed-accuracy trade-offs (*σ* ≈ 0), we regularized the fits by adding an additional term to the overall log-likelihood that penalized values of *σ*_*A*_ and *σ*_*B*_ that deviated from some typical value *σ*_0_:

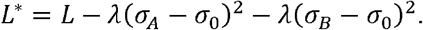

We set *σ*_0_ = 0.1*s* and *λ* = 500, which avoided overfitting the slope of the speed-accuracy trade-off without biasing parameter recovery too much. Our results were qualitatively unaffected by the exact values *σ*_0_ of and *λ* used.

## Acknowledgements

We thank E. Lesage and Y. Du for helpful comments on the manuscript, and M. Adputra for producing the stimuli. This project was supported by NSF Grant 1358756. RH is supported by Marie Skłodowska Curie Individual Fellowship NEURO-AGE (702784).

## Author Contributions

RH and AMH conceived and designed the experiments. RH collected the data. RH and AMH analyzed the data. RH and AMH wrote the manuscript. RH, AF, JK, and AMH reviewed and edited the manuscript.

## Materials & Correspondence

Corresponding Author: Robert Hardwick.

